# Harnessing Environmental DNA (eDNA) to Explore Frugivorous Interactions: A Case Study in Papaya (*Carica papaya*) and Pineapple (*Ananas comosus*)

**DOI:** 10.1101/2024.12.12.628216

**Authors:** Pritam Banerjee, Jyoti Prakash Maity, Nalonda Chatterjee, Sven Weber, Gobinda Dey, Raju Kumar Sharma, Chien-Yen Chen

## Abstract

Plant-animal interactions (PAIs) are critical in ecosystem function, mediating energy flow and species interactions. Traditional methods of tracking PAIs, such as morphological identification and camera trapping, are limited in speed and scalability, posing challenges for comprehensive biodiversity monitoring. Recently, environmental DNA (eDNA) metabarcoding has emerged as a promising technique for detecting species interactions non-destructively. This pilot study explores the application of eDNA metabarcoding to investigate interactions involving partially consumed and intact fruits of *Carica papaya* and *Ananas comosus*. eDNA metabarcoding were performed from of 36 partially consumed and 6 intact fruit samples. Metabarcoding of mitochondrial COI gene fragments revealed a diverse range of taxa, with Arthropoda, particularly insects, being the most abundant. Results indicated significant differences in taxonomic composition between pineapple and papaya samples, where both the fruit hold some unique as well as shared taxa. Furthermore, the diversity also differed between consumed and intact fruits, suggesting that partially consumed fruits serve as rich eDNA sources, capturing interactions with various frugivores and decomposers. Signal from various organisms detected through eDNA metabarcoding from consumed and damaged fruits allowed us to capture a wide array of taxa, revealing insights into species composition and ecological relationships. The unique ASVs associated with each fruit type suggest that certain taxa may showing preferences based on fruit characteristics such as sugar content, texture, or chemical profile. Present work highlighted the importance of eDNA based methods in unraveling the taxonomic composition of fruit-associated plant-animal interactions. This method needs limited taxonomic expertise, less labors, fast and effective, which can be implemented in monitoring ecological and economical species interactions.

## Introduction

Species interactions serve as significant conduits for energy transfer, with positive and negative interactions playing simultaneous roles in contributing to the ecosystem’s functioning (Åkesson et al., 2021). Plant-Animal Interactions (PAIs) are key drivers of energy flow and ecosystem functions, and understanding those interactions is the basis of understanding ecosystem functioning properties and processes. PAIs serve as a fundamental link between species, and those interactions are ecologically significant and can occur in various forms, including predation, frugivory, herbivory, parasitism, and mutualism, encompassing positive and antagonistic relationships (Banerjee et al., 2022). However, in the era of increasing anthropogenic activities (environmental pollution, climate change, species introduction, etc.), it’s getting difficult to trace the expeditious changes in our surroundings (Fletcher et al., 2024; Mazor et al., 2018; Cardinale et al., 2012), where the gradual loss of species interactions is leading to the extinction of many interconnected species (Valiente-Banuet et al., 2015). Maintaining species interaction balance in wildlife as well as agricultural ecosystems is essential. However, monitoring species diversity (native and/or introduce) and their interactions remains challenging. Traditional biomonitoring methods, such as morphological screening and camera trapping, are effective but lack the speed needed for timely detection (Banerjee et al., 2022). In the last few decades, the development of biomonitoring in different fields such as morphological screening with high-resolution microscopy, chemical taxonomy, radar-based remote sensing, automatic acoustic monitoring, and high-throughput sequencing methods for molecular monitoring are becoming the new standard across the globe (Van Van Klink et al., 2024; Chua et al., 2023). Especially, the recent development of metabarcoding and metagenomics allows us to deep-dive into environmental genomics, from presence/absence to abundance, species detection to species interactions, population genetics, and even gene expressions (Blackman et al., 2024; Stevens et al., 2023). More recently, environmental DNA (eDNA) based biodiversity monitoring gained solid attention due to its non-destructiveness, high accuracy, and cost-effectiveness over traditional methods. Thus, in the next generation of biodiversity monitoring, eDNA seems to be a promising technique for understanding spatial and temporal changes (Blackman et al., 2024). To date, eDNA has been successfully applied to answer many ecological questions such as presence and absence (Ficetola et al., 2008), relative abundance (Lacoursière-Roussel et al., 2016), habitat preference (Stoeckle et al., 2017), population genetics (Shum and Palumbi, 2021; Andres et al., 2023), and species interactions (Banerjee et al., 2022, Weber & Stothut et al. 2024). In some previous studies, traces of DNA recovered from flowers (Thomsen and Sigsgaard 2019; Johnson et al., 2023), leaves (Krehenwinkel et al., 2022), tree barks (Allen et al., 2023), honey (Utzeri et al., 2018) and even from tea (Krehenwinkel et al 2022) were used to understand the PAIs. Given the notable success of eDNA in understanding species interactions from different substrate, it can be hypothesized that fruits can be a potential source of eDNA to understand different label of frugivory. In a species-specific study, Monge et al. (2020) successfully amplified Macaw Bird’s DNA from partially consumed tropical almond fruits, which suggests the traces of DNA can be used to identify the interacting taxa. However, till now there are limited number of studies explored the understanding of frugivory interactions with eDNA metabarcoding. Metabarcoding of fruits can reveal many positive and negative interactions, and understanding those interactions are necessary in conservation biology, biosecurity and even in pest management agricultural ecosystem. Therefore, present study explored the application of fruit metabarcoding as a pilot study with two economically important fruits *Carica papaya* and *Ananas comosus*. Thus, we hypothesized that partially consumed or damaged fruits are a good source of eDNA for understanding the diverse interacting organisms. To test our hypothesis, we used partially consumed and unconsumed fruits of two economically important plants; *Carica papaya* and *Ananas comosus* from a locally grown agricultural farm near Chiayi County, Taiwan.

## Material and Methods

### Study area and sampling

Samples were collected from agricultural farms within a 5 km radius of National Chung Cheng University, Chiayi County, Taiwan. Two economically important fruiting plants, papaya (*Carica papaya*) and pineapple (*Ananas comosus*), were selected, and the fruits were divided into two categories: (i) partially consumed/damaged and (ii) intact without feeding marks. Due to difficulties in sampling partially consumed fruits, we conducted frequent farm visit between March and May 2022, collecting 18 partially consumed and 3 intact fruits per species. Each sample was collected with nitrile gloves in a separate sampling bag (Whirlpak®) and preserved in a cooler box with dry ice until returned to the laboratory. During our visit, we noted the presence of common Myna, Pallas’s squirrel, and many smaller animals like fruit flies. We also learned from communications with the farmers that the greater Bandicoot rat is a relatively common pest in these agricultural habitats.

### Accumulation of eDNA from fruits

Thirty-six partially consumed fruits and six intact fruits of papaya (*Carica papaya*) and pineapple (*Ananas comosus*) were processed for eDNA collection (Figure 1). To dislodge eDNA from the fruit surfaces, we sprayed the fruits with double-distilled water using a plastic spray bottle and collected the runoff in a 500 ml glass beaker. For each fruit, 300 ml of water was sprayed, and the collected water was filtered through a 0.45 μm pore-sized filter paper (GN-6 Metricel®, Pall Corporation) with a 47 mm diameter. After filtration, the filter papers were divided into two halves: one-half was preserved in Longmire’s buffer (Longmire et al., 1997), while the other half was stored in ATL buffer for subsequent DNA extraction.

**Figure 1.**
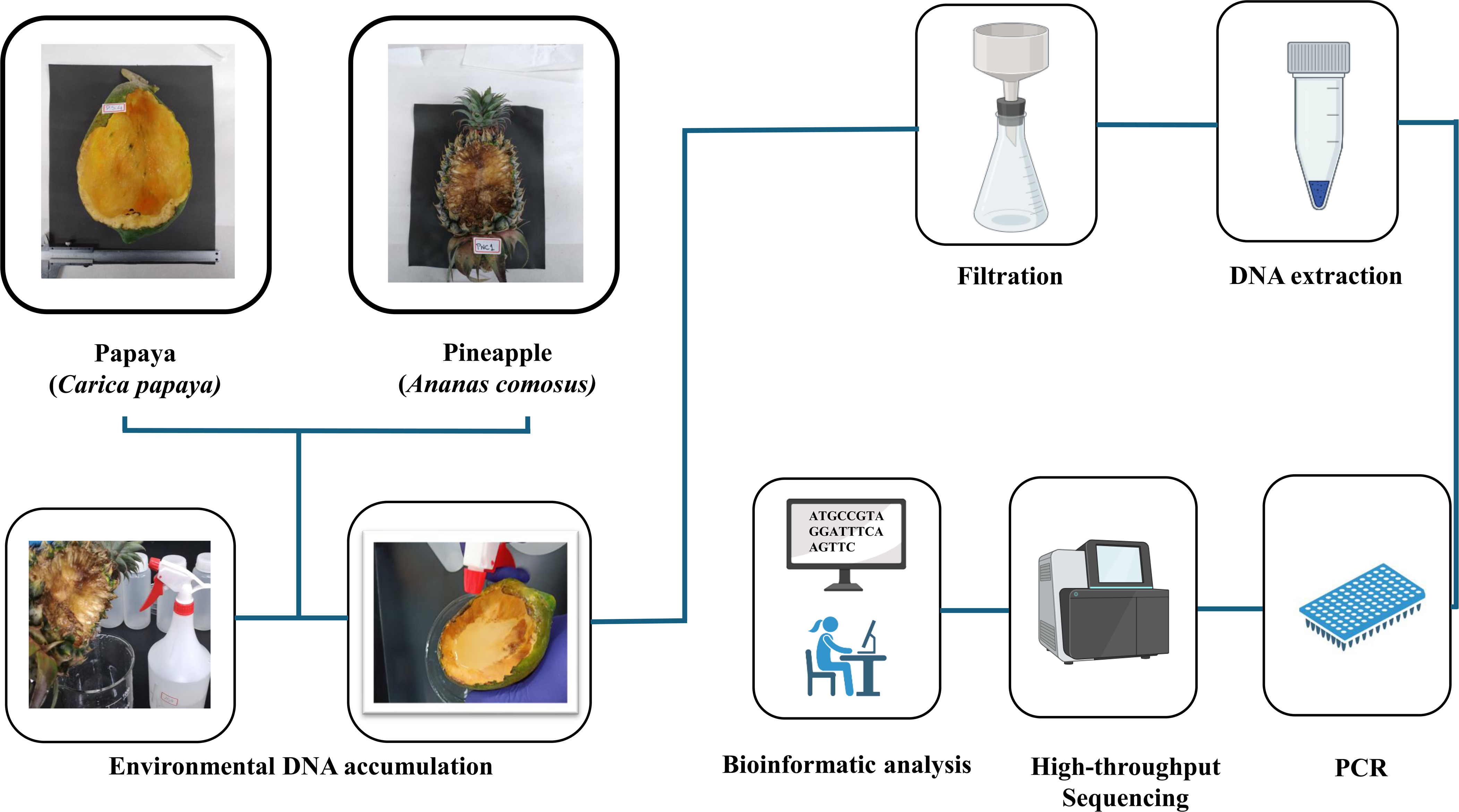
Diagrammatic workflow of environmental DNA (eDNA) metabarcoding from fruits of papaya (*Carica papaya*) and pineapple (*Ananas comosus*).

### DNA extraction and PCR amplification

We added 20 μl of Proteinase K to a tube containing 180 μl of ATL buffer and the filter paper. DNA extraction was performed using the DNeasy Blood & Tissue Kit (Qiagen, Germany) with the following modifications: (i) the filter paper was chopped into small pieces before lysis, (ii) lysis was carried out overnight at 60°C, and (iii) the spin column was incubated twice with 30 μl of AE buffer at 37°C for 10 minutes each. DNA quality and quantity were assessed using a Nanodrop spectrophotometer (Implen, Germany) and an Invitrogen Qubit 3.0 fluorometer (Thermo Fisher Scientific, USA). All samples, field blanks, capture blanks, and extraction blanks were extracted individually. Subsequently, we pooled three samples from the same or nearby plants (based on species) into a single composite sample, reducing the 18 partially consumed fruit samples to six samples (labeled as papaya: PP1–PP6 and pineapple: PN1–PN6) and the three intact fruits into one sample (labeled as papaya: PPC and pineapple: PNC) for each plant. These sample names were consistently used throughout the manuscript, including in figures and tables.

We amplified a portion of mitochondrial cytochrome c oxidase subunit I (COI) gene using universal degenerate primers: mICOIintF-5՛ -GGWACWGGWTGAACWGTWTAYCCYCC-3՛ and jgHCO2198-5՛ -TAIACYTCIGGRTGICCRAARAAYCA-3՛ specially designed for metazoans (Leray et al., 2013). Three PCR replicates were carried out for each sample (including blank samples) and the final reaction mixture of 25 μl was prepared by mixing 5 μl of 5x Fast-Run^TM^ Taq Master Mix with Dye (Protech Technology Enterprise, Taiwan), 0.5 μl each of 10 pmol forward and reverse primers (Genomics, Taiwan), 3 μl of the eDNA sample, and adding UltraPure™ DNase/RNase-Free Distilled Water (Thermo Fisher Scientific, USA) to make up the final volume. PCR was performed using a CLUBIO PCR system (CB series, Scientific Biotech Corp., Taiwan). The PCR protocol included an initial denaturation step at 94°C for 5 minutes, followed by 36 cycles of amplification: denaturation at 94°C for 30 seconds, annealing at 55°C for 30 seconds, and extension at 72°C for 1 minute. A final extension was performed at 72°C for 5 minutes. Each PCR was conducted in triplicate, and the subsamples were pooled based on band intensity observed during 1.5% agarose gel electrophoresis. The PCR products were then cleaned using 1.5x SPRI beads (AMPure XP Bead-Based Reagent) for library preparation.

### Library preparation and next-generation sequencing

The library was prepared using the TruSeq Nano DNA Library Prep Kit (Illumina, USA), and its quality was assessed with the Invitrogen Qubit 3.0 fluorometer (Thermo Fisher Scientific, USA) and the Agilent Bioanalyzer 2100 system (Agilent Technologies, USA). All qualified libraries were sequenced using Illumina MiSeq 300 bp paired-end reads (Illumina, USA) by Genomics Bioscience and Technology Co., Ltd., Taiwan.

### Data processing

The amplicon libraries were sequenced using the Illumina MiSeq platform (Genomics BioSci & Tech Co., New Taipei City, Taiwan). Paired-end reads (2 × 300 bp) were trimmed using Trimmomatic (version 0.39) (Bolger et al., 2014) and demultiplexed with an in-house script. Sequences from both ends of mlCOIintF and jgHCO2198 primers were trimmed using Cutadapt (version 3.4) (Martin, 2011) with the following criteria: read length ≥ 1 bp, minimum overlap ≥ 3 bp, and an error rate of 0.1 (default). Denoising, chimera removal, and assignment to Amplicon Sequence Variants (ASVs) were performed using DADA2 (Callahan et al., 2016). After error removal, all sequences were blasted against the GenBank nucleotide database in September 2022 using BLASTN, and taxa were selected based on the Lowest Common Ancestor (LCA) method (https://github.com/timkahlke/BASTA). The criteria for sequence alignment included 500 aligned sequences per query, with a 90% query cover and 80% to 100% sequence similarity. Reads with sequence similarity below 80% were excluded from the dataset. For taxonomic assignments, a similarity threshold of 90% was used for family-level, 95% for genus-level, and over 98% for species-level. Additionally, sequences assigned to genus and species were manually blasted against the NCBI database to confirm their identity.

### Statistical analysis

All statistical analyses were performed using R version 4.3.3 (R Core Team, 2020). A filtering procedure was applied to reduce the presence of artifacts, such as inflated read counts and false positives (see Ficetola et al., 2016). To visualize differences in taxonomic composition between the survey methods, a non-metric multidimensional scaling (NMDS) analysis was performed using Jaccard similarity matrices, with the vegan package in Rstudio (Oksanen et al., 2022). For general visualization the packages ggplot2 and ggvenn were used (Hamilto and Ferry, 2018). The significance of any observed differences between the fruits was tested using an Analysis of Similarity (ANOSIM) with Jaccard similarity matrices and 9,999 permutations. Additionally, to determine if there was a significant difference between the two fruits, we conducted a paired two-tailed t-test comparing the read counts per sample, both filtered and unfiltered. Rarefied ASV richness was calculated based on the minimum reads between all samples using the rarefy () function in vegan.

## Results

### Taxonomic composition

The Illumina MiSeq platform generated 1,692,642 single reads, corresponding to 796,234 paired reads, from 14 samples after filtering and denoising. On average, each fruit produced 56,873 ± 15,733 reads (Mean ± SD). All sequences were analyzed, and after database matching, amplicon sequence variants (ASVs) with identification below 80% were excluded. This resulted in the selection of 119 ASVs spanning diverse taxonomic groups, including metazoans, protozoans, algae, fungi, and some bacteria (supplementary_file_1). Given that universal degenerate primers amplify a broad range of taxa, reads from non-target groups (algae, fungi, and bacteria) were excluded, leaving 43 ASVs for analysis (supplementary_file_1). After classification, we identified 14 orders, 18 families, 15 genera, and 22 species. Among the total ASVs identified, the phylum Arthropoda was the most abundant (65.11%), followed by Rotifera (11.90%) and Chordata (9.50%) (supplementary_file_1). Within the phylum Arthropoda, the most common class was Insecta (71.43%), though other classes, such as Arachnida, Malacostraca, and Collembola, were also present (Figure 2, supplementary_file_1). At the genus level within Arthropoda, we identified three genera of fruit flies (*Drosophila* sp., *Zaprionus* sp., and *Bactrocera* sp.), two genera of beetles (*Araecerus fasciculatus* and *Carpophilus marginellus*), one genus of ants (*Tapinoma* sp.), parasitoid wasps (*Leptopilina* sp.), springtails (*Homidia sinensis*), isopods (*Burmoniscus* sp.), and non-biting midges (*Chironomus javanus*). Among the genus of fruit flies, we identified six species of *Drosophila*: *Drosophila takahashii*, *Drosophila bipectinata*, *Drosophila malerkotliana*, *Drosophila ananassae*, *Drosophila immigrans*, and *Drosophila eugracilis*, as well as one species of *Bactrocera*, *Bactrocera dorsalis*. Additionally, we identified one rodent species (the greater bandicoot rat, *Bandicota indica*), one squirrel species (Pallas’s squirrel, *Callosciurus erythraeus*), and one bird species (Javan myna, *Acridotheres javanicus*).

**Figure 2.**
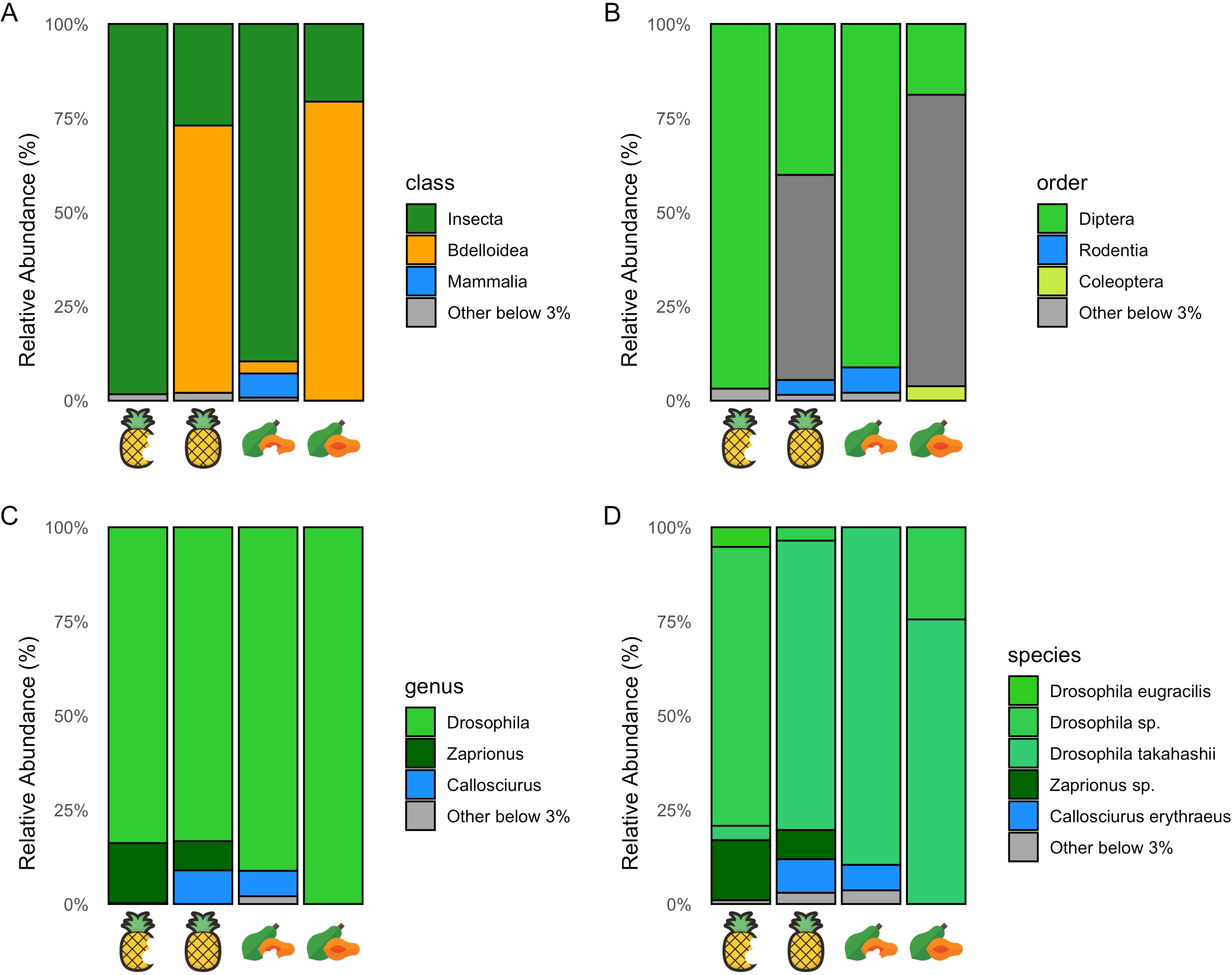
Top abundant taxa detected on consumed and intact fruits, categorized by (A) Class, (B) Order, (C) Genus, and (D) Species.

Among the 22 species, the fruit flies consist of the majority of the taxa, six species of *Drosophila*, one species of *Bactrocera,* and *Zaprionus* occupied most of the reads in our sample (Figure 2). However, we also detected other frugivores like *Acridotheres javanicus*, *Callosciurus erythraeus*. Furthermore, two species of beetle *Araecerus fasciculatus*, *Carpophilus marginellus* and one species of rat, *Bandicota indica* are serious crop pest. Furthermore, the genus *Leptopilina* sp., is known for *Drosophila* parasitoids.

### Consumed Pineapple vs Consumed papaya vs intact fruits

Comparison of alpha and beta diversity suggests that the taxonomic composition differs significantly between pineapple and papaya samples and within each of these sample types (Figures 3a and b). Furthermore, pineapple samples uniquely composed 15 ASVs, and papaya samples 11 ASVs, whereas together they share 16 ASVs (Figure 4). Most of the insect taxa were noted to be interacting with both the fruits, such as *Zaprionus* sp., *Drosophila takahashii*, *Drosophila eugracilis*, *Chironomus javanus*, *Callosciurus erythraeus* (Figure 5). However, some certain taxa noted to interact with either pineapple or papaya, such as *Drosophila ananassae*, *Drosophila bipectinata*, *Drosophila malerkotliana* interacting with pineapple samples, and *Tapinoma* sp., *Acridotheres javanicus*, *Araecerus fasciculatus*, *Burmoniscus* sp., were found to be interacting with papaya samples (Figure 5).

**Figure 3.**
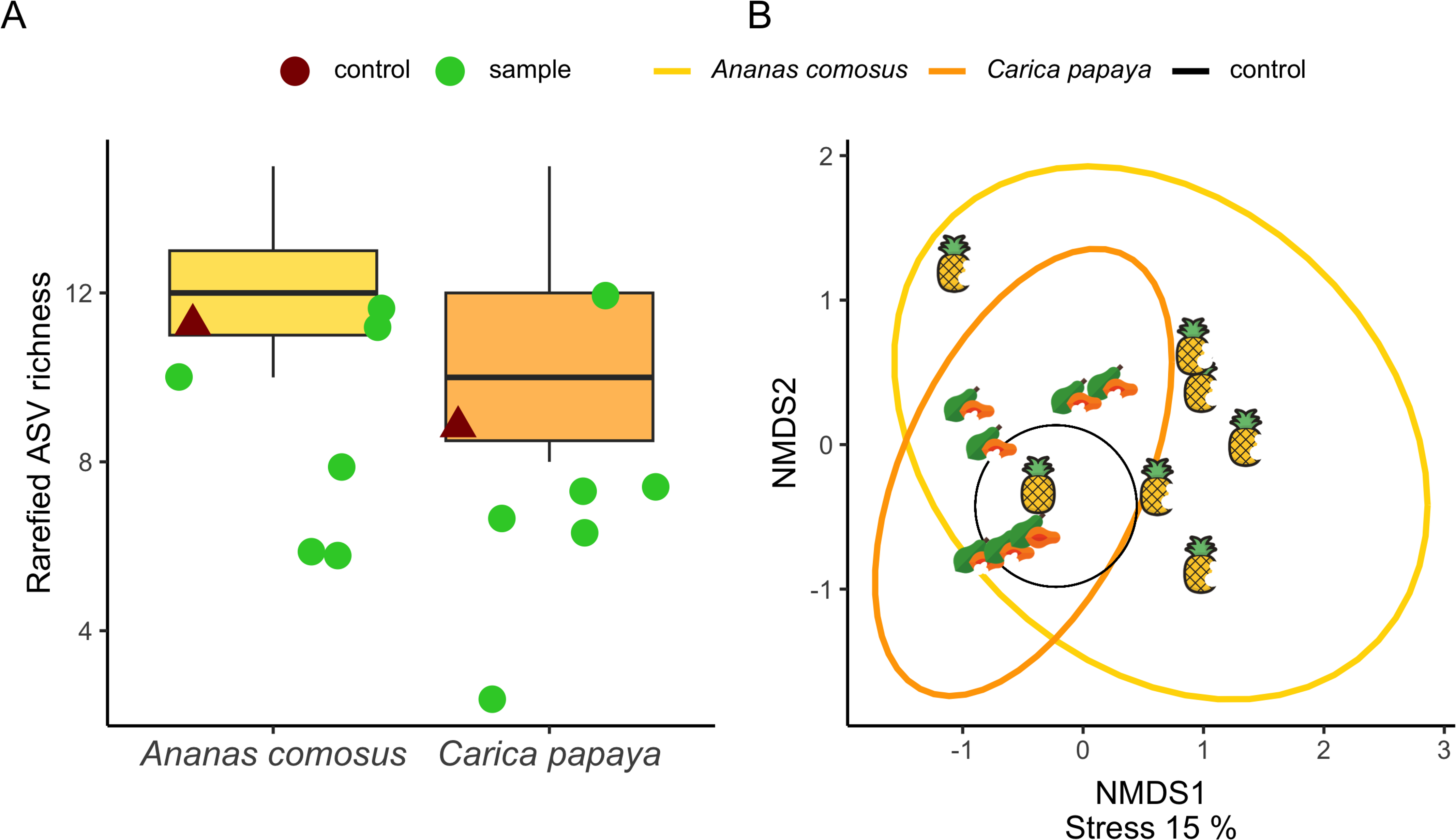
Sample orientation and richness from the fruits of papaya (Carica papaya) and pineapple (Ananas comosus), (A) Boxplot of ASV richness across consumed papaya (orange), consumed pineapple (yellow), and intact (uneaten) fruits (black).

**Figure 4.**
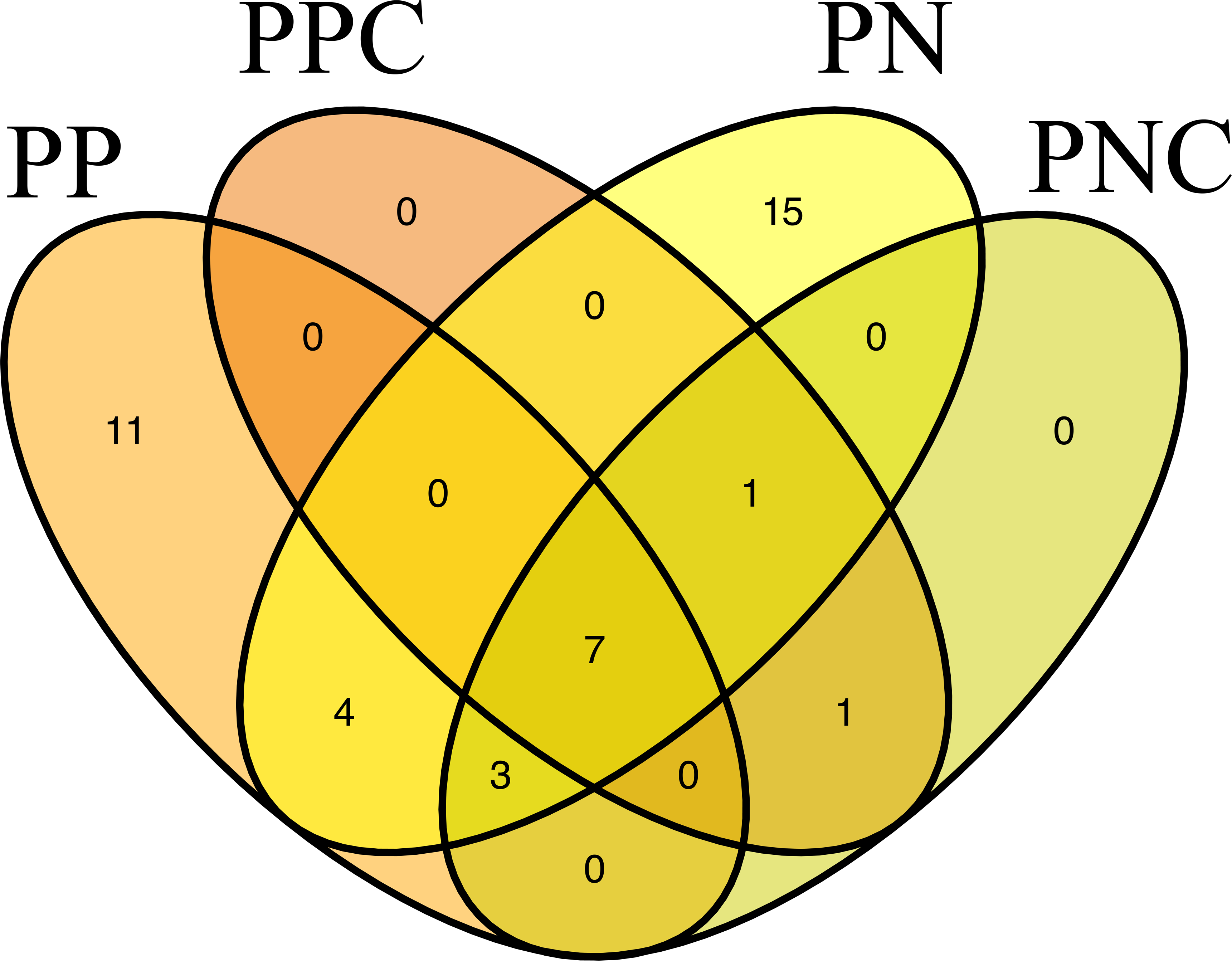
Heatmap of top 20 ASVs and their relative abundance in fruits of papaya (*Carica papaya*) and pineapple (Ananas comosus).

**Figure 5.**
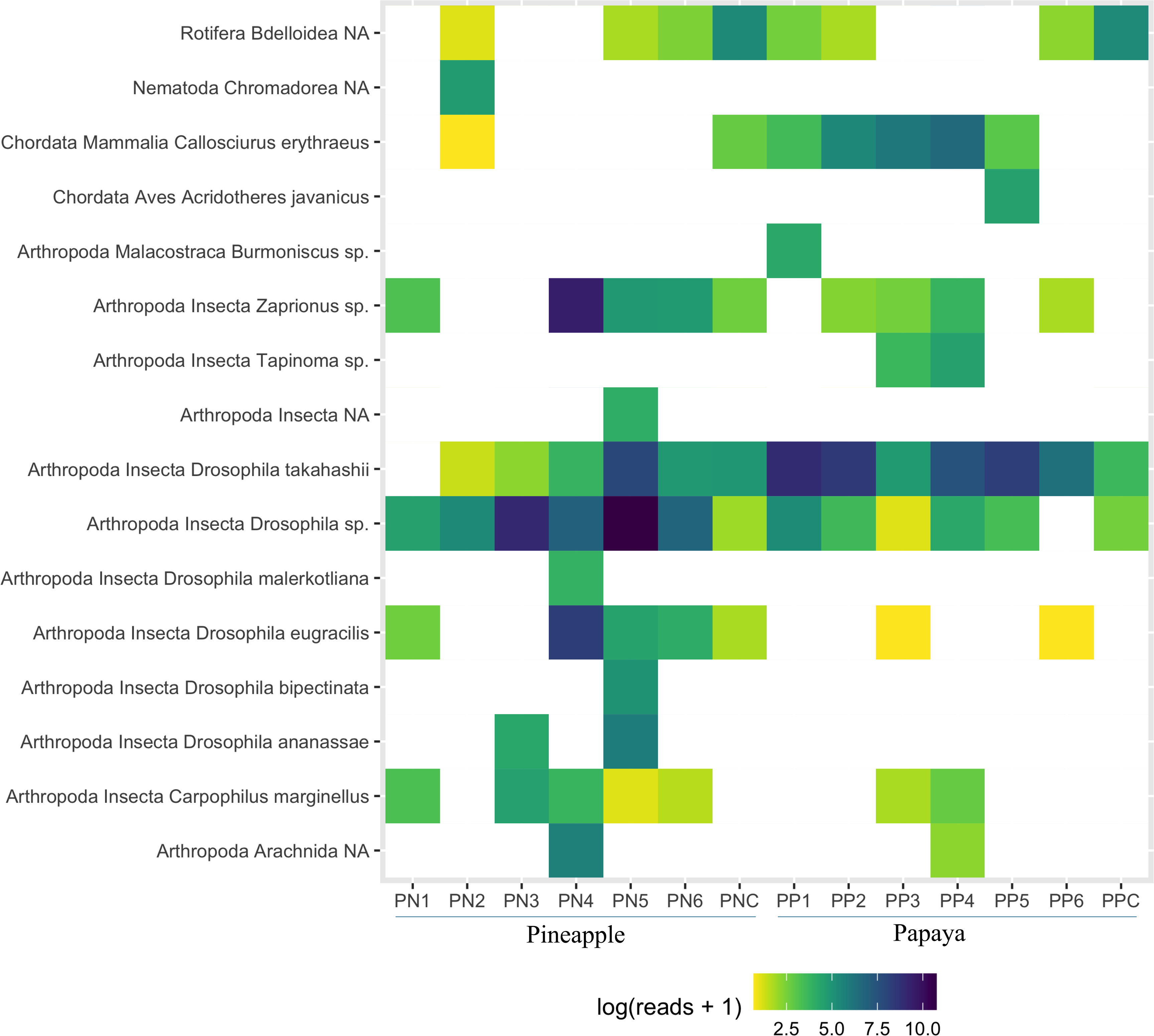
Ven diagram of overlapping ASVs between consumed fruits (PP: papaya; PN: pineapple) and intact fruits (PPC: papaya; PNC: pineapple).

The taxonomic composition of consumed and intact fruit significantly differed, as 15 ASVs and 11 ASVs were noted from consumed pineapple and papaya samples respectively, whereas one unique ASVs were reported in intact fruits (Figure 4). The overall ASV richness does not differ significantly between the consumed fruits. Furthermore, the ternary plot shows significant detection of arthropod families and orders towards consumed rather than intact fruits (Figure 6).

**Figure 6.**
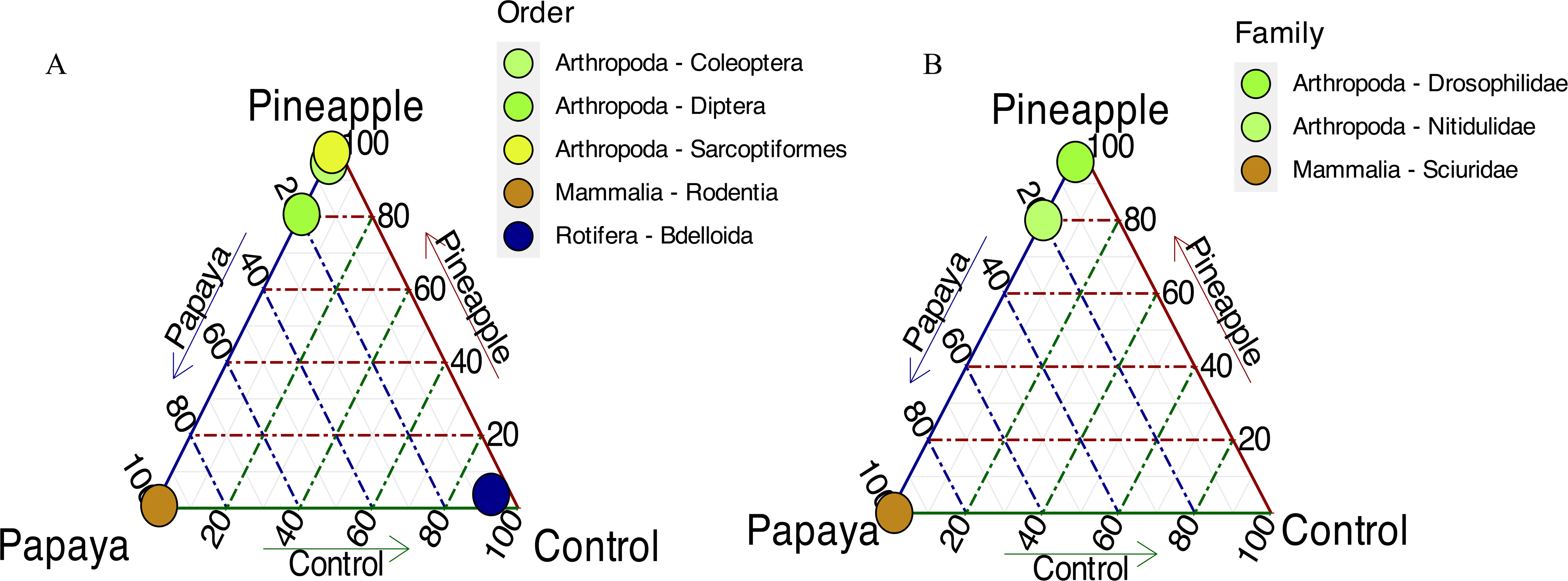
Ternary plot illustrating the distribution of individual ASVs in consumed pineapple (PP), consumed papaya (PP), and control samples (PPC and PNC), represented by (A) order and (B) family.

## Discussion

In this study, we successfully detected eDNA signature from various species interacting with fruits. Signal from various organisms detected through eDNA metabarcoding allowed us to capture a wide array of taxa in few sampling efforts, revealing insights into species composition and ecological relationships, particularly among insect species interacting with fruits. Traditionally those interactions were studied using morphological observation, and camera-trapping, however, our method could aid a clearer understanding of species interactions with less time and effort. Furthermore, eDNA methods could be better than individual barcoding or bulk-scale metabarcoding, where destructive measures need to be taken. Our results provide a comprehensive profile of the taxonomic diversity associated with fruits in both consumed and intact states, where consumed fruits were noted to have high traces of eDNA due to species interactions.

### Taxonomic Composition and Insect Community Structure

The taxonomic composition detected in our samples highlights the dominance of the phylum Arthropoda, particularly the class Insecta, which comprised the majority of ASVs, primarily from fruit-associated species (Figure 2, 5 and 6). The high proportion of fruit flies (*Drosophila* spp. and *Bactrocera dorsalis*) and beetles (*Araecerus fasciculatus* and *Carpophilus marginellus*) suggests these taxa are key components of the fruit ecosystem, playing likely roles as frugivores and decomposers (Deconninck et al., 2024). The detection of multiple *Drosophila* species aligns with previous studies that emphasize the adaptability of these flies to a range of fruit types and stages of decay (Deconninck et al., 2024). The presence of other arthropods such as springtails (*Homidia sinensis*), isopods (*Burmoniscus* sp.), and non-biting midges (*Chironomus javanus*) further reflects the complexity of fruit microhabitat. It suggests diverse ecological functions from detritivore to nutrient cycling within these communities.

Our findings of genera such as *Leptopilina*, a known parasitoid of *Drosophila*, underscore the presence of complex trophic interactions within the fruit environment. This suggests parasitoid-host dynamics, typically observed in natural settings, may also occur within these more transient microhabitats. This co-occurrence was also previously studied by researchers where those parasites were used as regular natural biological control of *Drosophila* (Chabert et al., 2012; Ibouh et al., 2019)

### Comparative Analysis of Pineapple and Papaya Communities

Our alpha and beta diversity analyses reveal significant differences in taxonomic composition between pineapple and papaya samples (Figure 3a, b). The unique ASVs associated with each fruit type suggest that certain taxa are specialized or may showing preferences based on fruit characteristics such as sugar content, texture, or chemical profile (Figure 4 and 5). For instance, species like *Drosophila ananassae* and *Drosophila malerkotliana* were primarily associated with pineapple, while other *Drosophila* species were predominantly found in both of the fruits. Also, taxa such as *Tapinoma* sp. (ants) and the introduced bird species, *Acridotheres javanicus* (Javan myna), were predominantly found in papaya samples. These findings suggest that specific ecological traits of the fruits may drive niche differentiation among taxa, leading to specialized interactions and potential competitive exclusion in these microhabitats. However, those traits need to be well studied on even in large scale to draw proper conclusion and connections among them.

### Differences Between Consumed and Intact Fruits

A notable difference in taxonomic composition was observed between consumed and intact fruits, with consumed fruits showing a higher ASV richness and diverse arthropod assemblages (Figure 2 and 4). This pattern likely reflects increased accessibility and nutrient availability due to open pericarp and ripening status of consumed fruits, which attracts a greater diversity of decomposers, frugivores, and opportunistic species. In contrast, the limited detection of ASVs in intact fruits, dominated by Rotifera reads, implies minimal ecological interaction, likely due to the intact physical barrier of pericarp and non-ripening status deterring colonization and minimizing DNA transfer, which proves our hypothesis.

The distinct arthropod associations with consumed fruits, as depicted in the ternary plot (Figure 6), underscore the attraction of these organisms to damaged or partially decayed fruits, which offer an enriched microenvironment compared to intact counterparts. This suggests open or decayed fruits attract a more complex community structure, facilitating ecological interactions across trophic levels. Here, eDNA can be very useful to study those ecological interactions precisely and timely. Furthermore, in agricultural systems, the presence of dropped or decayed fruits can potentially serve as a food source for introduce and/or pest species, thus eDNA can be a regular monitoring method to protect the economical damage in agricultural system.

### Implications for Invasion Ecology and Ecosystem Dynamics

The diverse array of detected species, including several introduced or pest species, highlights the role of fruits as critical interaction sites that may facilitate the establishment and spread of invasive species and pest species. For instance, *Bactrocera dorsalis* and *Acridotheres javanicus,* known agricultural pests, were identified within the fruit samples, indicating their capacity to exploit disturbed microhabitats. *Bactrocera dorsalis* is a highly invasive pest species around the Pacific-Asia Region that was first reported from Taiwan (Wan et al., 2012). This species is responsible for enormous economic loss in agriculture as their larvae feed on fruits and destroy the cultivation (Hardy et al., 1973; Wan et al., 2012). Additionally, the detection of species like *Bandicota indica*, a crop pest, suggests the potential for such interactions to impact surrounding agricultural ecosystems, leading to broader implications for pest management and biosecurity.

Overall, this study sheds light on the ecological complexity and taxonomic diversity associated with fruit habitats with eDNA metabarcoding, emphasizing the importance of monitoring fruit-associated communities for maintaining ecological balance and species interactions. This method also could be potentially useful for monitoring agricultural pest species in a short time and with limited effort. Future research should explore the temporal dynamics of these communities and assess how fruit decomposition stages further shape taxonomic composition. Also, exploring the species interactions study with eDNA for conservation and invasion biology can aid to proper management decision.

## Conclusion

Signal from various organisms detected through eDNA metabarcoding from consumed and damaged fruits allowed us to capture a wide array of taxa, revealing insights into species composition and ecological relationships. The unique ASVs associated with each fruit type suggest that certain taxa may showing preferences based on fruit characteristics such as sugar content, texture, or chemical profile. Present work highlighted the importance of eDNA based methods in unraveling the taxonomic composition of fruit-associated plant-animal interactions. This method needs limited taxonomic expertise, less labors, fast and effective, which can be implemented in monitoring ecological and economical species interactions. Moreover, this biomonitoring method can be used for regular monitoring assay in wildlife as well as agricultural field for suitable management decision.

## Acknowledgments

The authors would like to thank the Ministry of Science and Technology (Taiwan) for financial support (MOST 109-2811-M-194-502; MOST 108-2811-M-194-510).

## Data Availability Statement

Data sharing supporting the study will be openly available in dryad (https://doi.org/10.5061/dryad.k98sf7mhh), and its privately available for peer review here (Reviewer URL: http://datadryad.org/stash/share/AZa4n9v_p7NIrDzcJmkvNGZ36jLA4L769J1AP1kHHek.).

## Author contribution

P.B., J.P.M., and C.Y.C. designed the study. P.B., N.C., R.K.S., and G.D. performed the experiments. P.B. and S.W. conducted the data analysis. P.B. wrote the first draft of the manuscript. All authors provided extensive edits and contributed to the revision of the manuscript.

